# Disease mutations in the N-terminal extension and α2’-helix of dyskerin do not abolish dimerization but disrupt binding to the telomerase RNA

**DOI:** 10.1101/2025.08.22.671787

**Authors:** Jian Qin, Patrick Lambert-Lanteigne, Alexandre Garus, Chantal Autexier

## Abstract

The H/ACA ribonucleoprotein complex component dyskerin is essential for the biogenesis of H/ACA RNAs, including the human telomerase RNA (hTR). The N-terminal extension and α2’ helix of dyskerin are hotspots for disease-associated mutations linked to X-linked dyskeratosis congenita (X-DC), a premature aging disorder. Some of these mutations disrupt dyskerin–hTR interactions, leading to hTR destabilization and reduced telomerase activity. Cryo-EM structures of human telomerase have shown that the N-terminal extension and α2’ helix participate in dyskerin dimerization. However, biochemical evidence for dyskerin dimerization is still lacking, and it remains unclear whether mutations in these regions impair hTR binding by disrupting dimerization. Here, we provide the first biochemical evidence that dyskerin undergoes dimerization. We further demonstrate that dimerization is RNA independent and not abolished by disease mutations in the N-terminal extension or α2’ helix. Instead, these mutations impair hTR binding. Our findings offer new mechanistic insight into how mutations in the dyskerin N-terminal extension and α2’ helix contribute to the pathogenesis of X-DC.

## Introduction

Ends of linear chromosomes of eukaryotic organisms, known as telomeres, shorten after each DNA replication due to the end replication problem. Critically short telomeres trigger a DNA damage response, eventually leading to replicative senescence. To counteract telomere shortening, most eukaryotic organisms utilize telomerase to extend telomeres (1, 2). Telomerase is a holoenzyme comprised of the telomerase reverse transcriptase (TERT), the telomerase RNA (hTR in human) that contains the template to direct reverse transcription, and proteins involved in biogenesis of the telomerase RNA (3). Vertebrate telomerase RNAs have adopted the H/ACA domain at their 3’ end, which contains a 5ʹ hairpin followed by the H box (5ʹ ANANNA-3ʹ), a 3ʹ hairpin, and the ACA trinucleotide (4, 5). The H/ACA domain recruits dyskerin (Cbf5 in yeast), NHP2, NOP10, and NAF1 co-transcriptionally for hTR stabilization, with NAF1 later replaced by GAR1 to regulate biogenesis of the H/ACA ribonucleoprotein (H/ACA RNP) (3).

Mutations in dyskerin, NHP2, NOP10, and NAF1 have previously been found to cause hTR biogenesis defects and are associated with telomere maintenance disorders. Particularly, many dyskerin mutations are found in a premature aging disease known as X-linked dyskeratosis congenita (6–9). Dyskerin is a pseudouridine synthase that is highly conserved from archaea to eukaryotes to catalyze the conversion of uridine into pseudouridine in target RNAs. It is also crucial for the biogenesis and stabilization of RNAs containing the H/ACA domain (10). Human dyskerin consists of the catalytic domain and the C-terminal pseudouridine synthase and archaeosine transglycosylase RNA-binding domain (C-PUA) that are conserved between archaea and eukaryotes. A region upstream of the catalytic domain is spatially close to the C-PUA and is termed the N-terminal extended PUA domain (N-PUA), which is structurally conserved between archaea and eukaryotes (11, 12). In addition, the N- and C-terminal extension (NTE and CTE) are only conserved in eukaryotes (13) (Figure 1A).

**Figure 1.**
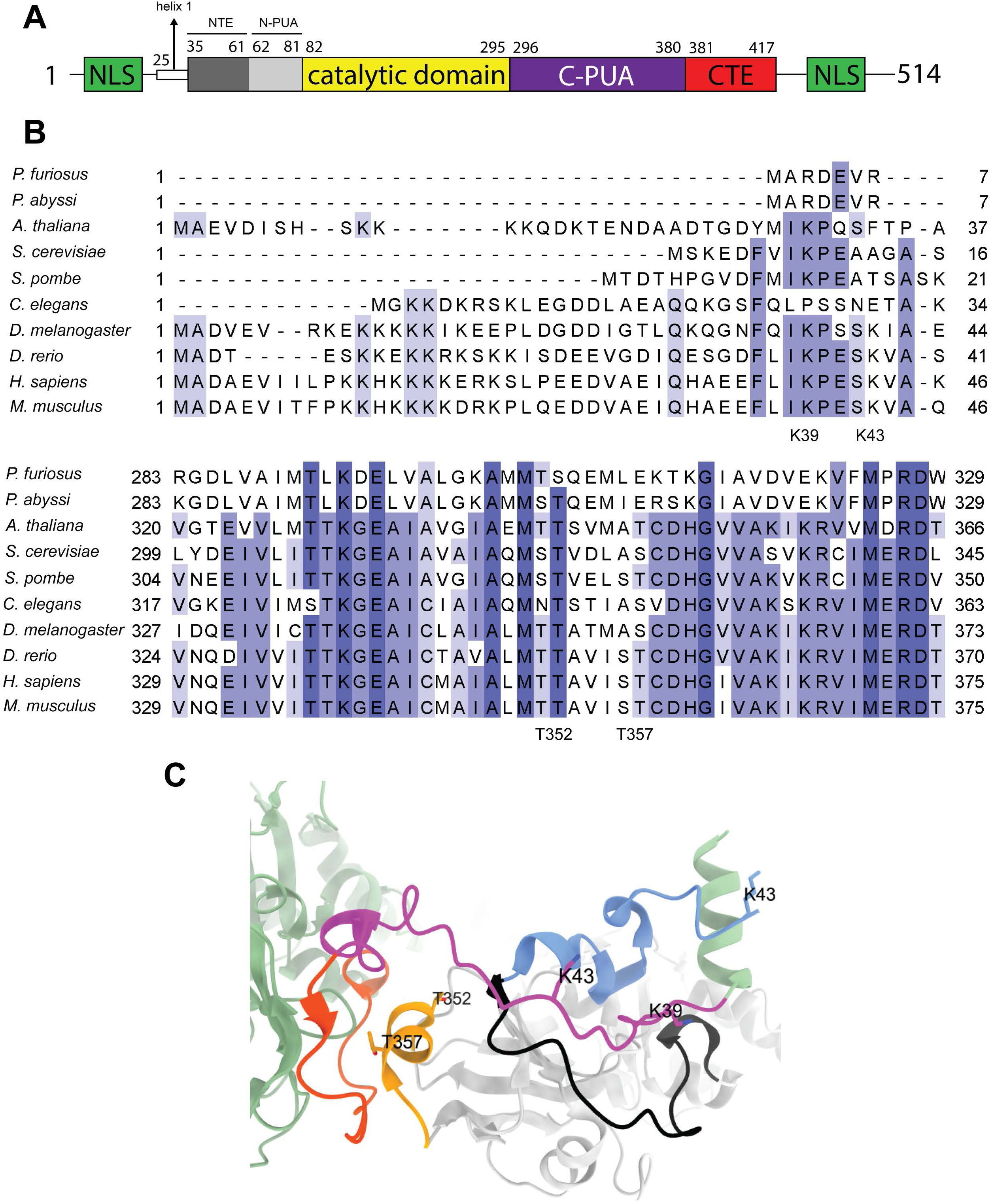
Schematic of human dyskerin, partial alignment, and cryo-EM structure of dyskerin in human telomerase. (A) Schematic of human dyskerin. NLS: nuclear localization signal; NTE: N-terminal extension; CTE: C-terminal extension. Helix 1 is resolved in 5’ dyskerin in a recent cryo-EM structure (15). (B) Partial alignment of human dyskerin with homologues in eukaryotes and archaea including the NTE (upper alignment) and α2’-helix (bottom alignment). The four disease-associated residues analyzed in this study are indicated. (C) cryo-EM structure of dyskerin in human telomerase (PDB: 8OUE) (15). Magenta: NTE of 5’ dyskerin; orange: N-PUA of 5’ dyskerin; green: the rest of the 5’ dyskerin; light blue: NTE of the 3’ dyskerin; black: N-PUA of the 3’ dyskerin; yellow: α2’-helix of the 3’ dyskerin; light grey: the rest of the 3’ dyskerin. Side chains of the four disease mutations studied in this article are shown. Other components of human telomerase are omitted for simplicity purposes.

Recent advances have been made towards understanding the structure of human telomerase and dyskerin. Within the telomerase holoenzyme, the two dyskerin molecules are engaged in intermolecular interactions (dyskerin bound to the 5’ and 3’ hairpin are referred to as 5’ and 3’ hereafter). A part of the NTE and N-PUA of the 3’ dyskerin form a hydrophobic pocket that contacts the NTE of the 5’ dyskerin; in return, a hydrophobic pocket in the 5’ dyskerin formed by the same regions as in the 3’ dyskerin interacts with the α2’-helix covering amino acids 350-360 in the C-PUA region of the 3’ dyskerin (14) (Figure 1B, C). A recent cryo-electron microscopy (cryo-EM) structure of human telomerase further resolved helix 1 (amino acids 25 - 33) upstream of the NTE in the 5’ dyskerin and amino acids 43 - 47 in the NTE of the 3’ dyskerin. These two regions form an additional interaction interface, where residues from helix 1 of the 5’ dyskerin interact with residues in the NTE and N-PUA of the 3’ dyskerin (15). Notably, the 5’ hairpin of hTR is only contacted by dyskerin, while the 3’ hairpin interacts with dyskerin, NOP10, NHP2 (16), and the association between the 3’ hairpin and the 3’ H/ACA protein complex is further enhanced by direct interaction between 3’ NHP2 and the biogenesis promoting box (BIO box) sequence on the 3’ hairpin (14). It is therefore proposed that the intermolecular interaction between the two dyskerin molecules compensates for the weak protein-RNA interaction at the 5’ hairpin of hTR by bridging the 5’ H/ACA protein complex to its 3’ counterpart that has strong interaction with the 3’ hairpin (16).

Consistent with the importance of the NTE, N-PUA, and the α2’-helix in mediating dyskerin dimerization, these three regions are mutational hotspots for X-DC. We have previously shown that K39E, K43E in the NTE as well as A353V in the α2’-helix reduced dyskerin binding to hTR (7). However, the mechanisms causing such defects are still not understood, as defects in dyskerin binding to hTR could be the result of impaired dimerization, or vice versa. In addition, previous studies assessing the interaction between dyskerin mutants and hTR in cells were performed without downregulating endogenous dyskerin, a scenario that does not recapitulate the disease situation where all dyskerin molecules carry the same mutation. Such an approach was initially adopted to avoid drastic reduction of total hTR levels that occurs upon dyskerin knockdown, in order to enable analysis of dyskerin binding to hTR by qPCR (6, 7, 17). However, lack of endogenous dyskerin knockdown creates a challenge interpreting results for mutants that are predicted to have defective dyskerin-dyskerin interaction: when these mutants show weakened association with hTR, it is unclear whether the defect is indirectly caused by the failure to interact with endogenous dyskerin rather than directly due to defects in dyskerin-hTR interactions. In this study, we first generated cell lines that allow inducible degradation of endogenous dyskerin. We then assessed the effects of K39E, K43E and two previously uncharacterized disease mutations in the α2’-helix, T352A and T357A, on dimerization and the interaction with hTR in the absence of endogenous dyskerin. Our study provides biochemical evidence for dyskerin dimerization consistent with the cryo-EM structures and demonstrates that the tested dyskerin mutants carrying disease mutations in the NTE and α2’-helix are defective in binding hTR but do not abolish dimerization, subcellular localization, or the interaction with other H/ACA RNP components.

## Results

### Generation of HCT116 dyskerin-DHFR cell lines

In previous studies we used siRNA to deplete endogenous dyskerin (6, 7, 17), since dyskerin knockout is embryonic lethal in mice (18). More recently, fusion of different degron systems with endogenous essential proteins by genome editing have allowed transient and inducible degradation of these proteins, omitting the needs for siRNA transfection (19, 20). We therefore inserted the *E. coli* dihydrofolate reductase (DHFR) degron downstream of dyskerin by CRISPR-Cas9-mediated knock-in. The dyskerin-DHFR fusion protein is spontaneously degraded, unless 10 μM of trimethoprim (TMP) is present. Due to the DKC1 gene encoding dyskerin being on the X-chromosome, we used HCT116, a colon cancer cell line derived from a male patient, to increase editing efficiency. From the six clones that survived antibiotic selection after editing, two have the endogenous dyskerin correctly fused to DHFR at the expected molecular weight (Figure 2). These two clones are referred to as C2 and C5 hereafter. PCR genotyping indicates that C2 might have a deletion downstream of the sequence corresponding to the 3’ homology arm in the repair template, while C5 might have an insertion in the sequence corresponding to the 3’ homology arm (Supplementary Figure 1). Nevertheless, our results suggest that in both clones, the fusion protein is at the expected molecular weight and can be degraded when TMP is removed for 24 h (Figure 2). These cell lines allow us to simultaneously express exogenous dyskerin and deplete endogenous dyskerin.

**Figure 2.**
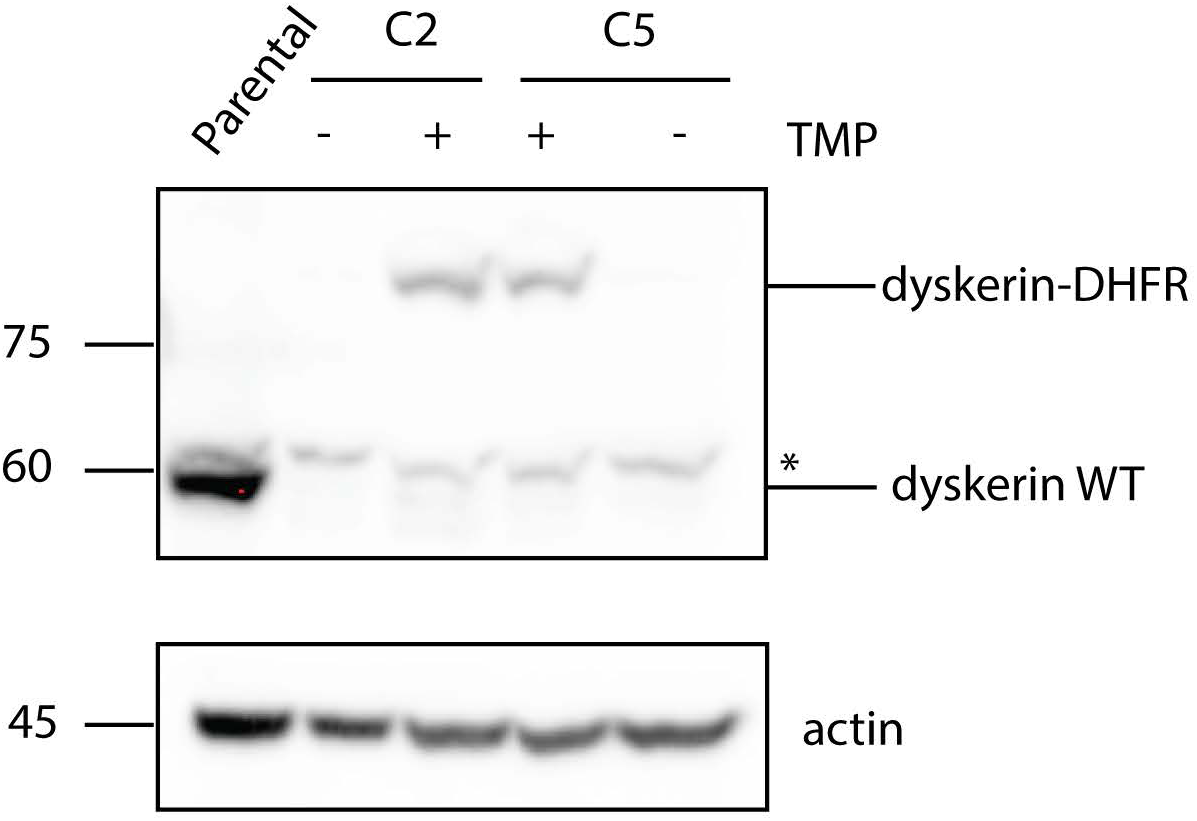
Verification of DHFR knock-in in HCT116 cells by western blot. Parental: HCT116 parental cells. C2 and C5 are two clones that express dyskerin-DHFR.TMP +: cells were constantly cultured in 10 µM of TMP. TMP –: cells were cultured in media with 0.1% DMSO without TMP for 24 h. Asterisk indicates non-specific band.

### Disease mutations in the dyskerin α2’-helix reduce hTR levels

We have previously shown that disease mutations in the NTE and α2’-helix: K39E, K43E, and A353V, failed to maintain hTR levels in HEK293 cells upon depletion of endogenous dyskerin by siRNA (7). To verify if two other uncharacterized disease mutations in the α2’-helix, T352A and T357A, also cause similar defects, we transiently expressed FLAG-dyskerin wild type or variants carrying these mutations in HCT116 dyskerin-DHFR C2 and C5 cells. Degradation of endogenous dyskerin and expression of FLAG-dyskerin was confirmed by western blot (Figure 3A). hTR levels were measured by qPCR upon induced degradation of endogenous dyskerin. Since mutations or depletion of dyskerin cause 3’ to 5’ degradation of hTR (21, 22), we used a primer pair that amplifies nucleotides 323 - 430 of hTR (23) to confirm its integrity at the 3’ end. In both cell lines, expression of the two mutants led to significantly lower hTR levels compared to expression of wild type FLAG-dyskerin upon degradation of endogenous dyskerin (Figure 3B). This verifies that our system can be used to study defects in maintaining hTR levels caused by mutations in dyskerin.

**Figure 3.**
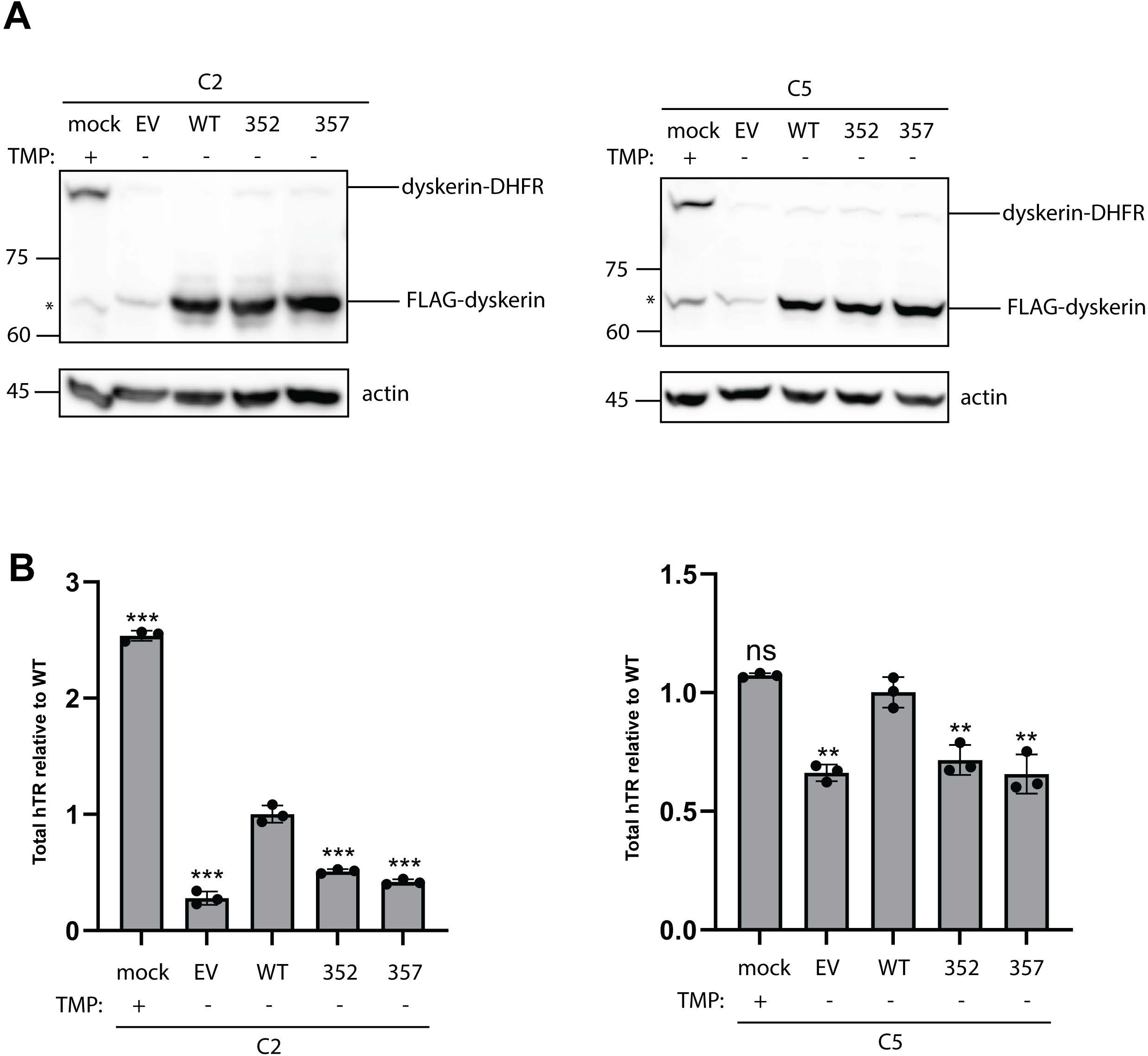
Dyskerin α2’ helix disease mutants T352A and T357A are defective in maintaining hTR levels upon induced degradation of endogenous dyskerin. (A) HCT116 dyskerin-DHFR C2 and C5 cells cultured in 10-cm plates were transfected with 10 µg of EV or FLAG-dyskerin. TMP was removed at the beginning of transfection for 48 h in total. Dyskerin protein levels were then analyzed by western blot using antibodies against dyskerin. Actin levels were assessed as a loading control. Asterisk indicates non-specific bands. Representative of three independent experiments is shown. (B) Total hTR levels in cells from (A) were measured by RT-qPCR and normalized to cells expressing wild type FLAG-dyskerin, using actin mRNA as the endogenous control. Bars represent the average values from three independent experiments; each data point indicates the value from one experiment. Error bars indicate standard deviation. **: P < 0.01; ***: P < 0.001. ns: statistically not significant.

### Disease mutations in the dyskerin dimerization interface are not sufficient to abolish dimerization

Recent cryo-EM structures of human telomerase have revealed intermolecular interactions between the two dyskerin molecules, which involves the NTE and N-PUA from amino acids 35-81, and the α2’ helix (amino acids 350 – 360) (14, 15) (Figure 1). We first tested whether dyskerin dimerization can be detected by immunoprecipitation and whether the dimerization is RNA-dependent in HCT116 dyskerin-DHFR C2 cells transiently transfected with FLAG- and HA-dyskerin. TMP was removed to degrade endogenous dyskerin. We show that pulldown of wild type FLAG-dyskerin also co-purified wild type HA-dyskerin and this interaction is not abrogated by RNase A/T1 digestion (Figure 4A). We then tested whether disease mutations in FLAG-dyskerin abolish the interaction with wild type HA-dyskerin. In this setting, FLAG-dyskerin and HA-dyskerin compete for binding to HA-dyskerin. Mutations in FLAG-dyskerin might cause it to be outcompeted by HA-dyskerin for binding to HA-dyskerin, which would be measured as a defect in dimerization. T352A and T357A as well as two previously characterized mutations in the NTE, K39E and K43E, were selected for the analysis. These single mutations in FLAG-dyskerin did not abolish the RNA-independent interaction with HA-dyskerin (Figure 4A). Next, we tested whether these mutations in both FLAG- and HA-dyskerin can diminish dimerization. This recapitulates the situation in patients where all dyskerin molecules carry the same mutation. We compared the amount of HA-dyskerin co-purified by anti-FLAG pulldown from cells transfected with wild type FLAG-dyskerin and HA-dyskerin mutant to those from cells transfect with FLAG- and HA-dyskerin carrying the same mutation. These experiments were also performed after induced degradation of endogenous dyskerin and in the presence of RNase A/T1. We found that wild type and all mutants of FLAG-dyskerin can readily interact with the HA-dyskerin mutants even after RNase digestion. The same disease mutation in both FLAG- and HA-dyskerin did not drastically reduce dyskerin dimerization compared to the situation where only HA-dyskerin carries the mutation (Figure 4B). Similar results were obtained in HCT116 dyskerin-DHFR C5 cells (Supplementary Figure 2).

**Figure 4.**
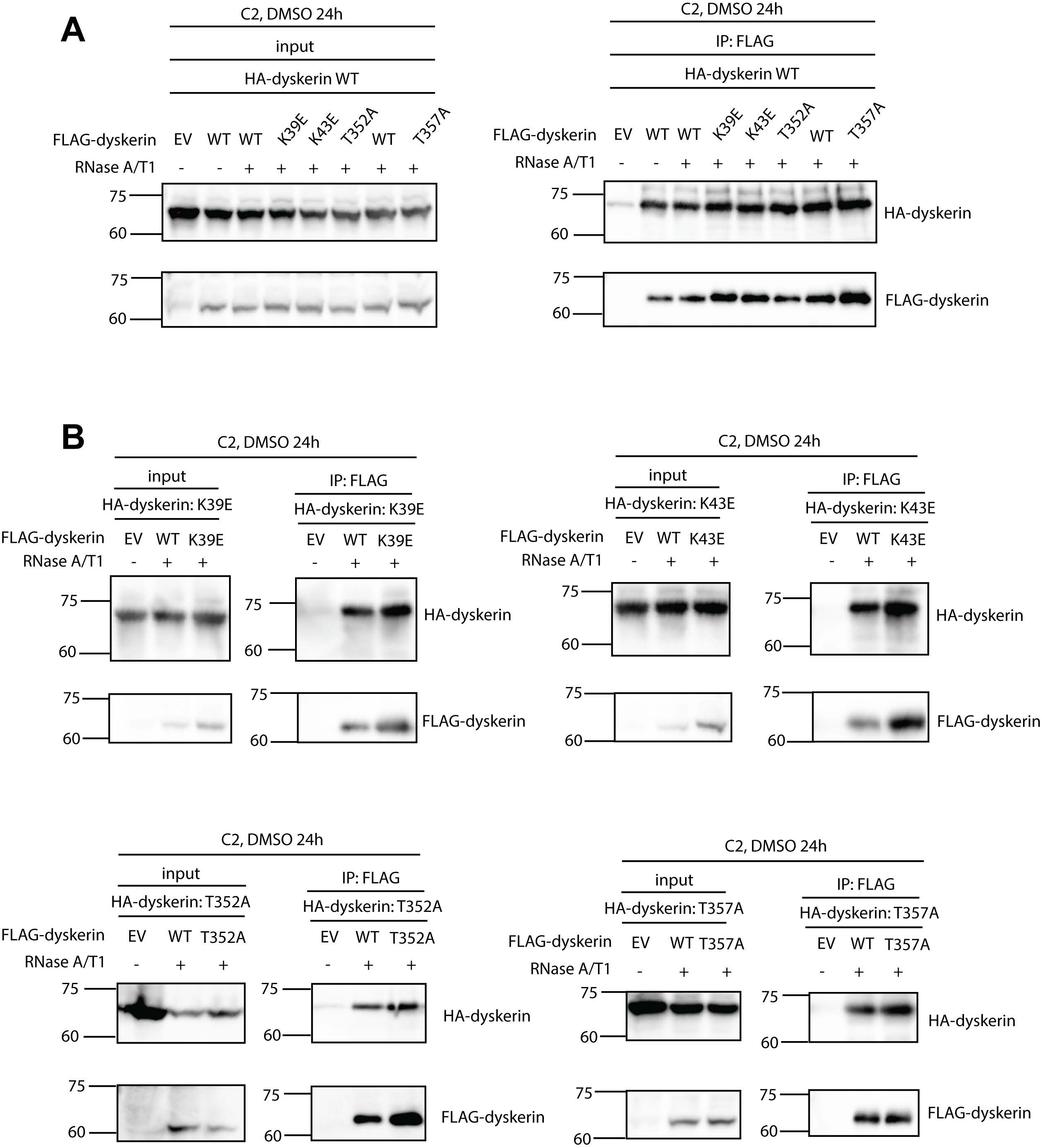
Disease mutations in the NTE and α2’ helix of dyskerin do not abolish the interaction between FLAG-dyskerin and HA-dyskerin. (A) (B) HCT116 dyskerin-DHFR C2 cells cultured in 10-cm plates were transiently transfected with 6 µg of EV/FLAG-dyskerin and 5 µg of HA-dyskerin. After removal of TMP for 24 h, immunoprecipitation using anti-FLAG magnetic beads was performed, followed by western blot against FLAG and HA. Presence or absence of RNaseA/T1 is indicated. Representative of two independent experiments is shown.

### FLAG-dyskerin carrying disease mutations in the dimerization interface are defective in binding to hTR in the absence of endogenous dyskerin

We previously analyzed dyskerin-hTR interaction without depleting endogenous dyskerin in order to maintain hTR levels in cells expressing dyskerin mutants, which is essential for qPCR analysis (6, 7, 17). Using our current system, we can simultaneously degrade endogenous dyskerin to prevents its interference and overexpress hTR so that cells transfected with the empty vector and FLAG-dyskerin wild type or variants have comparable levels of hTR. We performed RNA-IP in C2 and C5 cells co-transfected with 1 µg of hTR and 9 µg of empty vector or FLAG-dyskerin mutants (K39E, K43E, T352A, T357A) or co-transfected with 0.85 µg of hTR and 9 µg of FLAG-dyskerin wild type. Similar amount of FLAG-dyskerin wild type and mutants were pulled down (Supplementary Figure 3A; Figure 5A), and all cells had comparable total hTR levels (Supplementary Figure 3B; Figure 5B). K39E and K43E bound significantly less hTR than FLAG-dyskerin wild type in C2 cells (Supplementary Figure 3C), similar to the condition when endogenous dyskerin is present (24). Likewise, T352A and T357A showed significantly reduced interaction with hTR compared to wild type in both C2 and C5 upon TMP removal (Figure 5C). Together, our results showed that disease mutations in the dyskerin dimerization interface cause defects in hTR binding even in the absence of endogenous dyskerin.

**Figure 5.**
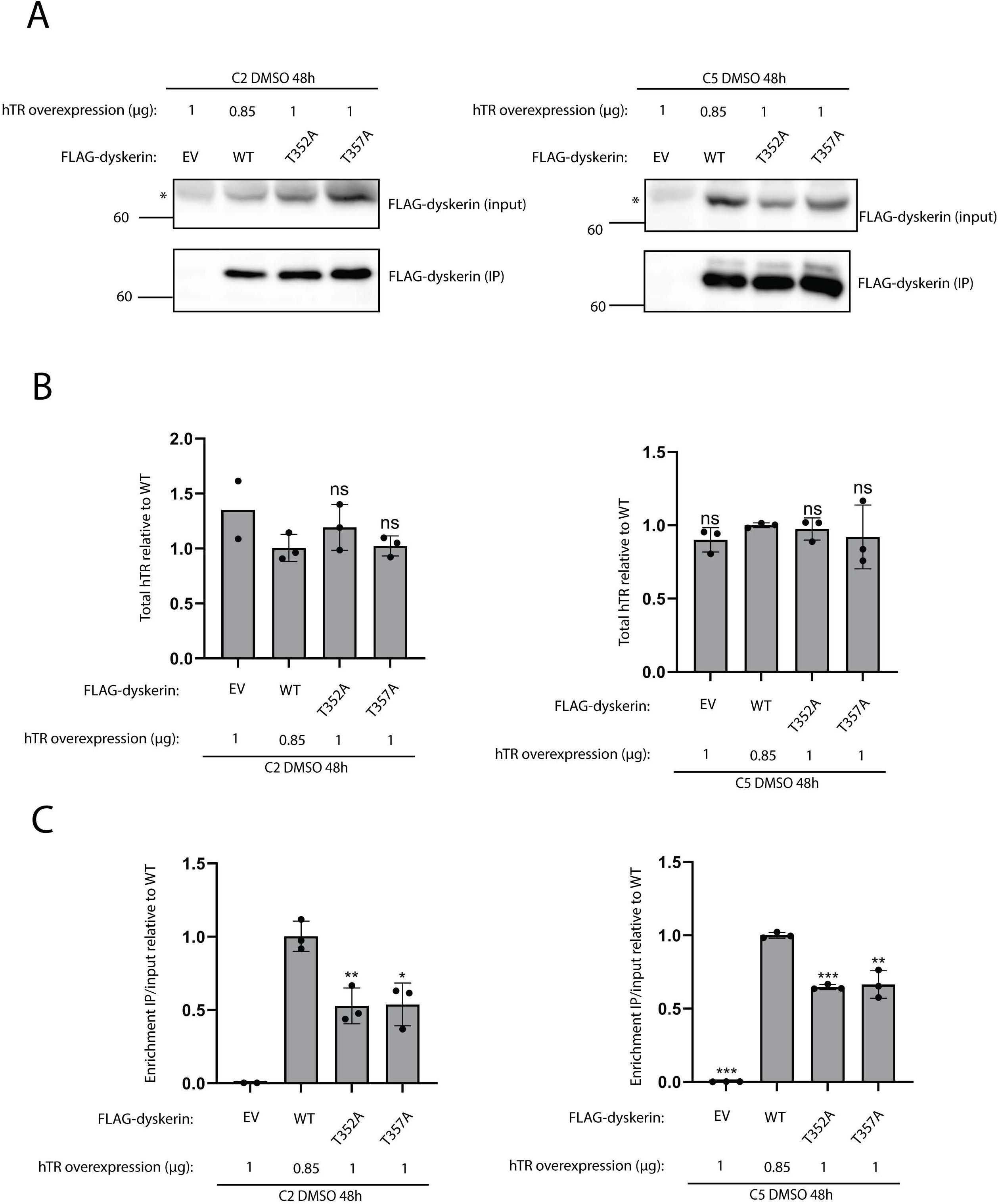
Dyskerin α2’ helix disease mutants T352A and T357A are defective in binding hTR upon induced degradation of endogenous dyskerin. (A) HCT116 dyskerin-DHFR C2 and C5 cells cultured in 10-cm plates were transiently transfected with 9 µg of EV/FLAG-dyskerin and the indicated amount of hTR. TMP was removed for 48 h. After immunoprecipitation with anti-FLAG magnetic beads, expression and pulldown of FLAG-dyskerin were verified by western blot using antibody against FLAG. Representative of three independent experiments is shown. (B) Total hTR levels in the input fraction from (A) were measured by RT-qPCR and normalized to cells expressing wild type FLAG-dyskerin, using actin mRNA as the endogenous control. Bars represent the average values from three independent experiments; each data point indicates the value from one experiment. Note that only two experiments for EV in C2 were performed. Error bars represent standard deviation. ns: statistically not significant. (C) Enrichment of hTR in the IP fraction over the input fraction was normalized to cells expressing wild type FLAG-dyskerin. Bars represent the average values from three independent experiments; each data point indicates the value from one experiment. Two experiments for EV in C2 were performed. Error bars represent standard deviation. *: P < 0.05; **: P < 0.01; ***: P < 0.001.

### Disease mutations in the dyskerin dimerization interface localize to Cajal bodies (CBs) and the nucleolus without abolishing the interaction with protein components of the H/ACA RNP

CBs and the nucleolus are two compartments essential for hTR processing and maturation, as well as for dyskerin-mediated pseudouridylation (10, 25). Defective localization to these compartments has been documented in DC or X-DC(6, 26). To assess whether the tested mutations also impair localization, we stained FLAG-dyskerin together with the CB marker coilin or with the nucleolar marker fibrillarin in dyskerin-DHFR C2 and C5 cells (Figure 6A; Figure 7A; Supplementary Figures 4 and 5) following degradation of endogenous dyskerin (Figure 6B; Figure 7B; Supplementary Figures 4 and 5). The K39E and K43E variants showed no defects in CB or nucleolar localization (Figure 6C; Figure 7C; Supplementary Figures 4 and 5), consistent with previous findings in HEK293 cells with endogenous dyskerin present (7). Similarly, T352A and T357A did not display any localization defects to both compartments (Figure 6C; Figure 7C; Supplementary Figures 4 and 5).

**Figure 6.**
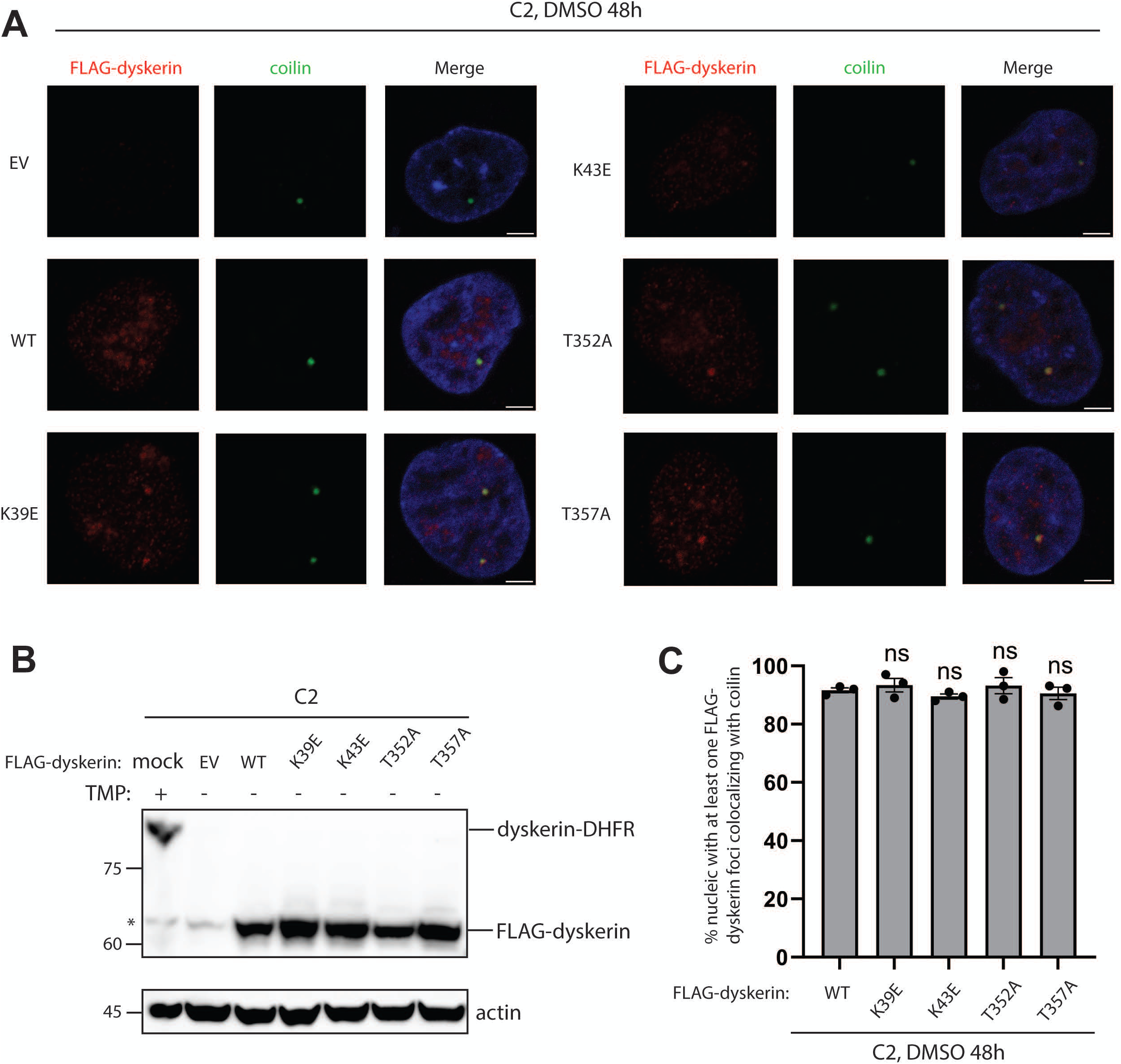
Disease mutations in the NTE and α2’ helix of dyskerin do not disrupt localization of dyskerin to CBs. HCT116 dyskerin-DHFR C2 cells cultured in 6-cm plates were transiently transfected with 6 µg of EV/FLAG-dyskerin in TMP-free media for 6 h. Transfection mixture was then removed, and the cells were cultured in TMP-free media for another 42 h. (A) Dual indirect immunofluorescence was performed using primary antibodies against FLAG and the CB marker coilin. Nuclei were stained by DAPI. Scale bars indicate 5 µm. (B) Degradation of dyskerin-DHFR and expression of FLAG dyskerin was confirmed by western blot. Asterisk indicates non-specific band. (C) Quantification of the percentage of nuclei with at least one FLAG-dyskerin foci co-localizing with coilin. Bars represent the average values from three independent experiments; each data point indicates the value from one experiment. At least 50 nuclei positive for both FLAG and coilin staining were examined in each experiment. Error bars represent standard deviation. ns: statistically not significant.

**Figure 7.**
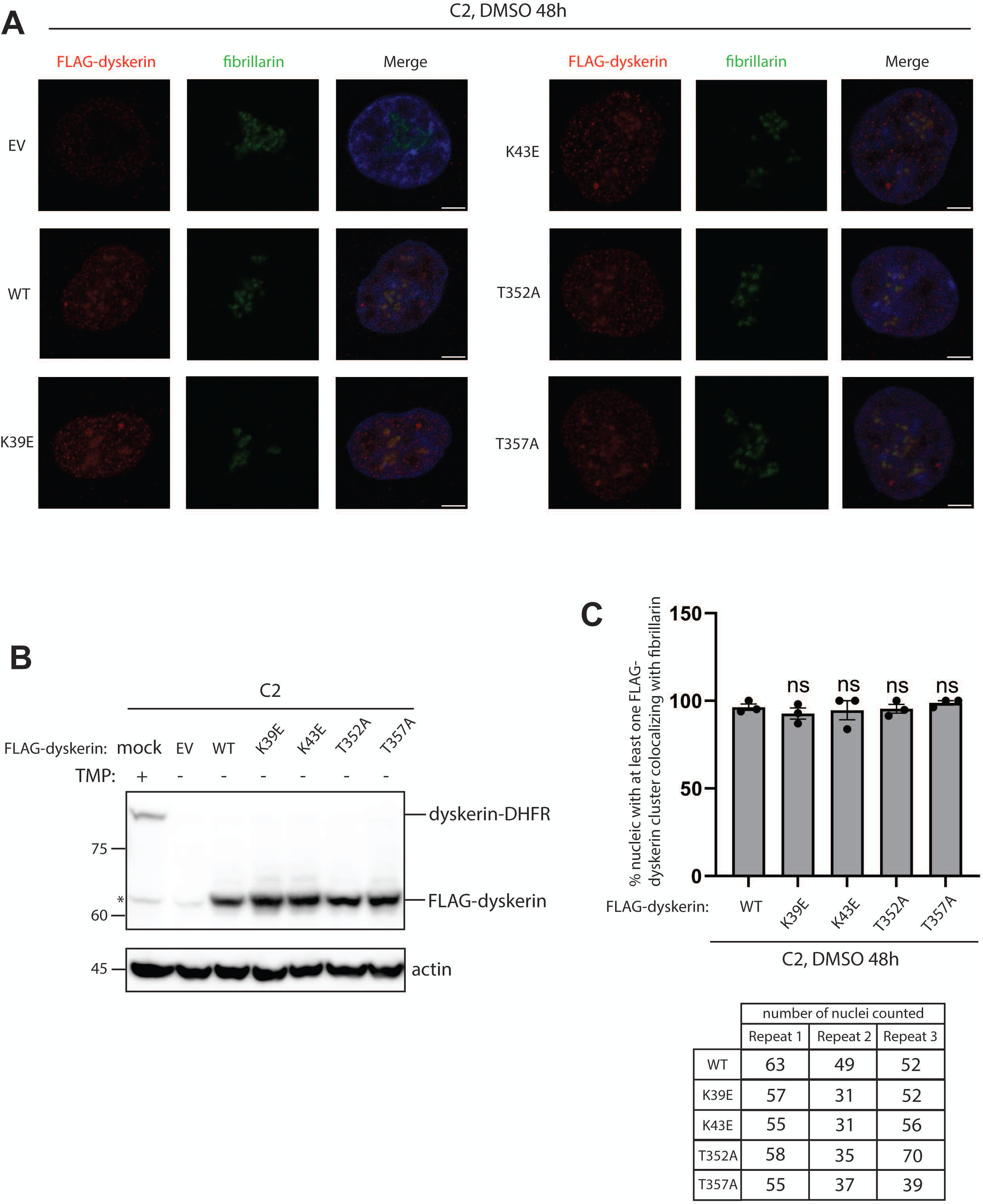
Disease mutations in the NTE and α2’ helix of dyskerin do not disrupt localization of dyskerin to the nucleolus. HCT116 dyskerin-DHFR C2 cells cultured in 6-cm plates were transiently transfected with 6 µg of EV/FLAG-dyskerin in TMP-free media for 6 h. Transfection mixture was then removed, and the cells were cultured in TMP-free media for another 42 h. (A) Dual indirect immunofluorescence was performed using primary antibodies against FLAG and the nucleolar marker fibrillarin. Nuclei were stained by DAPI. Scale bars indicate 5 µm. (B) Degradation of dyskerin-DHFR and expression of FLAG dyskerin was confirmed by western blot. Asterisk indicates non-specific band. (C) Quantification of the percentage of nuclei with at least one FLAG-dyskerin clustering co-localizing with fibrillarin. Bars represent the average values from three independent experiments; each data point indicates the value from one experiment. Error bars represent standard deviation. ns: statistically not significant. The number of nuclei examined, positive for both FLAG and fibrillarin staining, were listed in the table.

To examine whether the impaired hTR binding observed for these mutants was due to defective H/ACA RNP assembly, we immunoprecipitated FLAG-dyskerin wild-type and variants and analyzed their interactions with premature and mature H/ACA RNP components NAF1, NHP2, and GAR1. Since NOP10 bridges the interaction between dyskerin and NHP2 (27), NHP2 was used as an indirect readout for dyskerin-NOP10 interaction. Binding of the K39E and K43E variants to NAF1, NHP2, and GAR1 was not abolished, consistent with previous findings (7) (Figure 8AB). Similar results were obtained for T352A and T357A (Figure 8AB). Notably, in contrast to our previous findings in HEK293 cells where GAR1 appeared as a faint band between two non-specific bands in the IP fraction (6, 7, 17), here we observed a clear GAR1 band, possibly due to the use of different cell lines and the improvement of IP protocol. Taken together, the tested disease mutations in the dimerization interface are not sufficient to abolish correct nuclear localization or the interaction with H/ACA RNP components.

**Figure 8.**
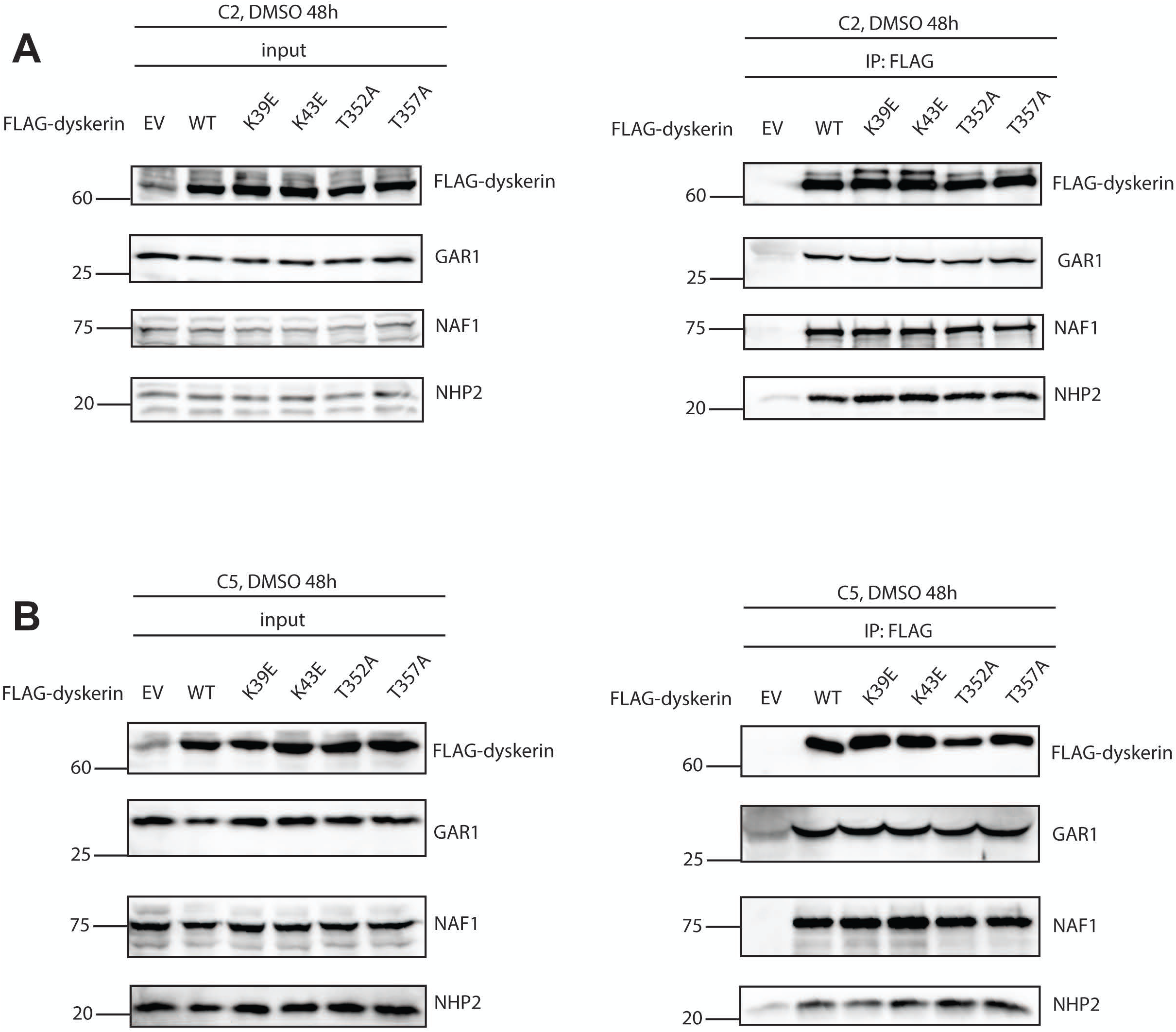
Disease mutations in the NTE and α2’ helix of dyskerin do not abolish the interaction between FLAG-dyskerin and HA-dyskerin. HCT116 dyskerin-DHFR C2 (A) and C5 (B) cells cultured in 10-cm plates were transiently transfected with 10 µg of EV/FLAG-dyskerin. After removal of TMP for 48 h, immunoprecipitation using anti-FLAG magnetic beads was performed, followed by western blot against FLAG, NAF1, GAR1, and NHP2. Three independent experiments were performed for C2 and two for C5.

## Discussion

Dyskerin plays an essential role in hTR biogenesis and telomerase regulation. The NTE and α2’ helix are two hotspots that contain multiple mutations associated with X-DC (7). While many mutations in the α2’ helix remain uncharacterized, how mutations in the NTE affects interaction with protein binding partners and hTR as well as subcellular localization has only been examined in the presence of endogenous dyskerin (24). Recent discoveries from cryo-EM structures showing the involvement of the NTE and α2’ helix in dyskerin dimerization (14, 15) emphasize the need for a system that alleviates the influence of endogenous dyskerin in the above processes. In this study, we generated two independent clones of HCT116 dyskerin-DHFR cells that allow inducible degradation of endogenous dyskerin by removing TMP from the culture media. We first verified that hTR levels decreased in both clones upon TMP removal, which was only rescued by expression of FLAG-dyskerin wild type but not EV or the mutants. Note that in C5, expression of FLAG-dyskerin wild type fully restored hTR to the level displayed by untransfected cells cultured with TMP, whereas the rescue was only partial in C2. This is possibly attributed to clonal differences. Nevertheless, the mutants were defective in maintaining hTR levels in both C2 and C5.

We provided biochemical evidence of dyskerin dimerization for the first time using immunoprecipitation and further showed that this interaction is not abolished by RNase treatment. The observation that dyskerin dimerization is resistant to RNase digestion suggests that dimers of dyskerin and the H/ACA protein complexes might form prior to dyskerin binding to H/ACA RNAs, and that interaction between the two dyskerin molecules might not only occur in telomerase but also in other H/ACA RNPs. This is consistent with a previous study, in which H/ACA protein complexes purified by pulldown of NHP2 were able to form RNPs with either the 3’ hairpin alone or both the 5’ and 3’ hairpins of the ACA28 snoRNA (28). Dyskerin dimerization is not abolished by disease mutations in the dimerization interface, not even when both FLAG- and HA-dyskerin carry the same mutation. We reasoned that mutating one residue might not be sufficient to completely abrogate such dimerization. Complete abolishment of dyskerin dimerization might require deletion of both the NTE and N-PUA.

Due to the unconventional structure of its 5’ hairpin, it has been speculated that hTR might only recruit one copy of the H/ACA protein complex to the 3’ hairpin. The presence of two copies of the protein complex was first shown by a biochemical analysis (29) then confirmed by the first cryo-EM structure of human telomerase (16). The same structure showed that the 5’ hairpin of hTR is only contacted by dyskerin, while the 3’ hairpin is bound by dyskerin, NHP2 and NOP10. This implies that the 5’ H/ACA protein complex might require an extra anchor to remain bound to hTR. Cryo-EM structures of human telomerase with higher resolution have revealed interactions between the two H/ACA protein complexes involving the 5’ and 3’ dyskerin and the 3’ NOP10 (14, 15), suggesting that optimal binding of dyskerin to hTR requires both direct dyskerin-hTR interaction and dyskerin dimerization. We previously showed that K39E, K43E, and A353V reduced dyskerin binding to hTR when endogenous dyskerin was present. In this study, we further demonstrated that K39E, K43E, T352A, and T357A are defective in binding hTR even in the absence of endogenous dyskerin. Since these mutants do not abolish dimerization, we hypothesize that at least some of these mutations directly disrupt dyskerin-hTR interaction, which is consistent with the 5’ NTE being in proximity to nucleotides in the ACA box of hTR (15). Alternatively, these mutations may still cause subtle dimerization defects not detectable by immunoprecipitation or western blot, but nonetheless sufficient to impair RNA binding.

Most of the H/ACA RNAs in archaea contain only one hairpin followed by the ACA box which is bound by a single set of Cbf5, Nop10, L7ae, and Gar1 (30). This raises the possibility that eukaryotic Cbf5/dyskerin might also possess the intrinsic ability to bind to a single hairpin in H/ACA RNAs. Supporting this idea, a previous study has shown that pulldown of dyskerin co-purified the single 3’ hairpin but not the single 5’ hairpin of hTR (28), consistent with the weakened interaction of the 5’ hairpin of hTR with the H/ACA protein complex (16). To clarify the mechanisms underlying hTR binding defects caused by mutations in the dimerization interface, it would be informative to test the binding of dyskerin dimerization mutants to a single 3′ hairpin of hTR in the future. Such an approach could help distinguish whether the observed hTR binding defects arise from direct impairment of dyskerin–hTR interaction or indirectly from compromised dyskerin dimerization.

## Experimental procedures

### Plasmids

The pcDNA3.1 FLAG-dyskerin plasmid was generated by sub-cloning FLAG-dyskerin into the HindIII site of pcDNA3.1 (-) neo, using pcDNA3.1-dyskerin (gift from Dr François Dragon, Université du Québec à Montréal) as a template for PCR amplification (31, 32). pcDNA3.1 3×HA-dyskerin is a gift from Dr. Steven Artandi (Standford University). pcDNA3.1 FLAG-dyskerin and pcDNA3.1 3×HA-dyskerin were used to generate all disease mutants by site-directed mutagenesis. All mutations were verified by Sanger sequencing covering the entire FLAG-dyskerin region by Genome Quebec. pBluescript II SK(+)-hTR is a gift from Dr. Joachim Lingner (École Polytechnique Fédérale de Lausanne) (33).

### Generation of HCT116 dyskerin-DHFR cell lines by CRISPR-Cas9-mediated knock-in

The human colon cancer cells HCT116 (gift from Dr. Alain Nepveu, McGill University) in a 24-well dish at about 50% confluency were transfected with recombinant TrueCut Cas9 v2 nuclease (Invitrogen) and TrueGuide Modified sgRNA (Invitrogen, sequence in Supplementary Table 1) using Lipofectamine CRISPRMAX (Invitrogen) according to the manufacture’s instruction. 300 ng of gel-purified dsDNA repair template was co-transfected. The repair template was generated by PCR amplifying the region in the pAc5 HA3-eDHFR-T2A-puro plasmid (Addgene, plasmid #86395) containing the sequences of genes encoding 3×HA-tagged *E. coli* dihydrofolate reductase (DHFR), a T2A self-cleavage peptide, followed by a puromycin resistance gene, and the SV40 polyadenylation signal. The primers used for amplification (purchased from IDT) contain overhangs of genomic DNA sequences of dyskerin flanking the stop codon (Supplementary Table 1). 10 μM TMP and 1 µM of Alt-R HDR enhancer v2 (IDT) were added to the media together with the transfection mixture. 24 h later, the media was changed to DMEM with 10% FBS, Antibiotic-Antimycotic, 10 μM TMP. After 72 h of initial transfection, cells were trypsinized, well resuspended to separate single cells, transferred to a 10 cm dish, and constant pressure of 1 μg/ml puromycin was applied. Colonies that survived the selection were manually selected and cultured individually. Insertion of DHFR and degradation of the fusion protein in the absence of TMP was confirmed by Western blot. PCR genotyping using Platinum™ Direct PCR Universal Master Mix (Invitrogen) was performed to detect aberrant insertion or deletion (primer sequences in Supplementary Table 1).

### Cell culture and transfection

HCT116 parental cells were cultured in DMEM with 10% FBS (Wisent) and Antibiotic-Antimycotic (Gibco). HCT116 dyskerin-DHFR cells were cultured in DMEM with 10% FBS, Antibiotic-Antimycotic, 1 μg/ml puromycin, and 10 μM trimethoprim (TMP) dissolved in DMSO. When TMP is not included in the culture media, DMSO at a final concentration of 0.1% is added instead. All cells were routinely tested to be free of mycoplasma contamination using the mycoplasma PCR detection kit (ABM, cat. G238).

Transient transfection of HCT116 cells was performed using Lipofectamine 3000 (Invitrogen) according to the manufacturer’s instructions. For Figures 3, 5, 6, and Supplementary Figure 2, the cells were kept in DMEM with 10% FBS, 0.1% DMSO, and the transfection mixture for 48 h; For Figure 5, the cells were grown in DMEM with 10% FBS and 10 μM TMP with the transfection mixture for 24 h, then in DMEM with 10% FBS and 0.1% DMSO for another 24 h; For Figures 6, 7, and Supplementary Figures 4, 5, the cells were kept in in DMEM with 10% FBS and 10 μM TMP with the transfection mixture for 6 h, then in DMEM with 10% FBS and 0.1% DMSO for another 42 h.

### Immunoprecipitation

To examine the interaction between FLAG-dyskerin and HA-dyskerin, endogenous NHP2, NAF1, and GAR1, cells were lysed in lysis buffer (25 mM HEPES, 150 mM KCl, 1.5 mM MgCl_2_, 10% glycerol, 0.5% NP-40, 5 mM β-mercaptoethanol, 1×protease inhibitor cocktail) and kept on ice for 30 min. Lysate containing 3 mg of proteins was incubated with 40 μl of 25% slurry Pierce Anti-FLAG Magnetic Agarose (Thermo Scientific, A36797) for 2 h at 4 °C. The beads were then washed four times with 1 ml of the corresponding lysis buffer. When indicated, 40 μg/ml (200 U/ml) of RNase A and 500 U/ml of RNase T1 were added to the 4^th^ wash. In these experiments, samples with and without RNase were incubated at 37 °C with rotation for 1h. RNase digestion was performed under more stringent treatment conditions than previously used to determine that PABPN1 interacts with dyskerin in a RNA-dependent manner (23), including higher amounts of RNases (200 U vs 7.5 U RNase A, 500 U vs 300 U RNase T1), a more optimal temperature for RNase activity (37 °C vs 25 °C), and longer duration of treatment (1 h vs 30 min). For all experiments, the washed beads were resuspended in 2×Laemmli buffer (120 mM Tris-HCl pH6.8, 20% glycerol, 4% SDS, 0.02% bromophenolblue, 5% β-mercaptoethanol). Lysate containing 150 μg of proteins was mixed with the same volume of 2×Laemmli buffer and kept as input.

### Whole cell lysate preparation and immunoblotting

To extract whole cell lysate, cell pellets were lysed in WCE buffer (20 mM HEPES-KOH pH7.9, 25% glycerol, 0.01 mM EDTA pH8.0, 0.2% NP-40, 5 mM MgCl_2_, 420 mM KCl, 1×protease inhibitor cocktail, 1 mM NaF, 1 mM NaVO_4_, 50 mM Na-β-glycerophosphate) on ice for 10min, then centrifuged at 15800 g for 10 min at 4 °C. After measuring the amount of proteins with Bradford assay, the supernatant was mixed with the same volume of 2×Laemmli buffer.

Whole cell lysate or protein samples from immunoprecipitation were resolved by SDS-PAGE, transferred to PVDF membrane, blocked in 1% BSA in PBS 0.1% Tween, incubated with primary and secondary anti-bodies, developed in ECL Plus Western Blotting Substrate (Thermo Scientific), and scanned in a ChemiDoc MP Imaging System (Bio-Rad). Primary antibodies used are anti-FLAG (Proteintech, 66008-4-Ig, 1:1000), anti-NAF1 (Abcam, ab157106, 1:1500), anti-NHP2 (Proteintech, 15128-1-AP, 1:2000), anti-dyskerin (Cell Signaling Technology, D6N4K, 1:2000), anti-actin (Cell Signaling Technology, 8H10D10, 1:10000), anti-GAR1 (Proteintech,11711-1-AP, 1:2000), anti-HA (Cell Signaling, C29F4, 1:2000).

### RNA-IP

Cells were lysed in RNA-IP buffer (25 mM HEPES, 150 mM KCl, 1.5 mM MgCl2, 10% glycerol, 0.5% NP-40, 1 mM DTT, 40 U/ml RNaseOUT, 1×protease inhibitor cocktail) and kept on ice for 30 min. Lysate containing 3 mg of proteins was incubated with 40 μl of 25% slurry Pierce Anti-FLAG Magnetic Agarose (pre-blocked with 10 μg/ml of yeast tRNA) for 2 h at 4 °C. The beads were washed four times with lysis buffer. One fraction of input lysate containing 150 μg of proteins and 5 μl of the beads was mixed with the same volume of 2×Laemmli buffer. Another fraction of input and the rest of the beads were resuspended in TRIzol (Life Technologies) for RNA extraction.

### RNA extraction and qPCR

Cell pellets, cell lysate or beads resuspended in Trizol were proceeded to RNA extraction according to the manufacturer’s instructions. The RNA pellets were resuspended in sterile water and treated with Turbo DNA-free kit (Life Technologies).

RNA samples were reverse transcribed into cDNA with random primers (ThermoScientific) and SuperScript II reverse transcriptase (Invitrogen) or with the qScript cDNA SuperMix (Quantabio) according to the manufacturer’s instructions. Reverse transcription products were diluted 10 times and subjected to qPCR analysis (5 μl SYBR-Green [Quantabio], 0.4 μl of 10 μM primer mix, 2.6 μl sterile H2O, 2 μl diluted reverse transcription product). The reactions were performed using the 7500 Fast Real-Time PCR system (Life Technologies), managed by the Applied Biosystems 7500 Software v2.3. The reaction condition was as followed: 50 °C for 20s, 95 °C for 10 min, 40 cycles of (95 °C for 15 s and 60 °C for 1 min). To measure total hTR levels, actin mRNA was used as the endogenous control. For RNA-IP analyses, the corresponding RNAs in the input fraction were used as the endogenous control. Fold change of each sample relative to the control was calculated according to the methods in (34). Primer sequences used in qPCR are provided in Supplementary Table 1.

### Immunofluorescence

Cells grown on cover slips were rinsed with PBS, fixed in PBS-buffered 4% formaldehyde for 10 min at room temperature, washed in PBS three times for 5 min each, followed by permeabilization in PBS-buffered 0.5% TritonX-100 for 10 min at room temperature. Permeabilized cells were washed in PBS three times for 5 min each, then blocked in PBG (1% v/v fish gelatin [Sigma, G-7765], 0.5% w/v BSA in PBS) for 30 min at room temperature. To assess localization of FLAG-dyskerin to Cajal bodies, cells were incubated with rabbit anti-FLAG (Sigma-Aldrich F7425, 1:500) and mouse anti-coilin (gift from Dr. Michael Terns, University of Georgia, 1:10000) in PBG overnight at 4 °C. To assess localization of FLAG-dyskerin to the nucleolus, cells were incubated with rabbit anti-FLAG and mouse anti-fibrillarin (monoclonal antibody 72B9 obtained from Dr. Kenneth Michael Pollard, The Scripps Research Institute (35), 1:30) in PBG overnight at 4 °C. After primary antibody incubation, cells were washed in PBG three times for 5 min each, followed by incubation with secondary antibodies (FITC-donkey anti-mouse IgG, Jackson Immuno Research Lab Inc., 1:125; or Cy3-donkey anti-rabbit, Jackson Immuno Research Lab Inc., 1:125) in PBG for 1 h at room temperature. Cover slips were then washed in PBG twice for 5 min each, rinsed in PBS, and mounted in Vectashield with DAPI (Vector Laboratories) (6). Images were captured by LSM800 confocal microscope (Zeiss) provided by the Core Facility at the Lady Davis Institute, at 63×magnification using the ZEN Pro software.

### Multiple sequence alignment

Clustal Omega (36) was used to generate multiple sequence alignment of human dyskerin, its homologues in *Pyrococcus furiosus*, *Pyrococcus abyssi*, *Arabidopsis thaliana*, *Saccharomyces cerevisiae*, *Schizosaccharomyces pombe*, *Caenorhabditis elegans*, *Drosophila melanogaster*, *Danio rerio*, *Mus musculus*. Amino acid sequences of each protein were obtained from UniProt (identifiers O60832 for human, Q7LWY0 for *P. furiosus*, Q9V1A5 for *P. abyssi*, Q9LD90 for *A. thaliana*, P33322 for *S. cerevisiae*, O14007 for *S. pombe*, O17919 for *C. elegans*, O44081 for *D. melanogaster*, F1Q749 for *D. rerio*, Q9ESX5 for *M musculus*).

### Statistical analyses

Statistical analyses were performed in GraphPad Prism9. The control was compared to other samples using unpaired parametric t-test. P-value < 0.05 indicates statistical significance.

## Data availability

All data are contained within the manuscript. All raw data supporting the results of this study are available from the corresponding author on request.

## Supporting information

This article contains supporting information.

## Acknowledgements

We thank Dr François Dragon (Université du Québec à Montréal) for providing the pcDNA3.1-dyskerin plasmid; Dr. Steven Artandi (Standford University) for providing the pcDNA3.1 3xHA-dyskerin plasmid; Dr. Joachim Lingner (École Polytechnique Fédérale de Lausanne) for providing the pBluescript II SK(+)-hTR plasmid. We thank Dr Michael Terns (University of Georgia), Dr Kenneth Michael Pollard (The Scripps Research Institute) for providing mouse anti-coilin, and mouse anti-fibrillarin antibodies. We thank Dr. Alain Nepveu (McGill University) for providing HCT116 parental cells.

## Author contributions

J.Q. and C.A. conceptualization; J.Q. methodology; J.Q., P.L.-L., A.G. performed experiments; J.Q. writing-original manuscript; C.A. writing-review and editing; C.A. supervision and funding acquisition.

## Funding and additional information

Financial support was provided from Canadian Institute for Health Research Grant PJT-166130 to C.A. Support to J.Q. was provided by a McGill University Faculty of Medicine Internal Studentship (Max E. Binz Fellowship) and Fonds Recherche Quebec-Santé (FRQS) doctoral award. A.G. is supported by a McGill University Faculty of Medicine Internal Studentship (J.P. Collip Fellowship) and FRQS doctoral award.

