## Supplemental Figures for "Disease mutations in the N-terminal extension and α2’-helix of dyskerin do not abolish dimerization but disrupt binding to the telomerase RNA"

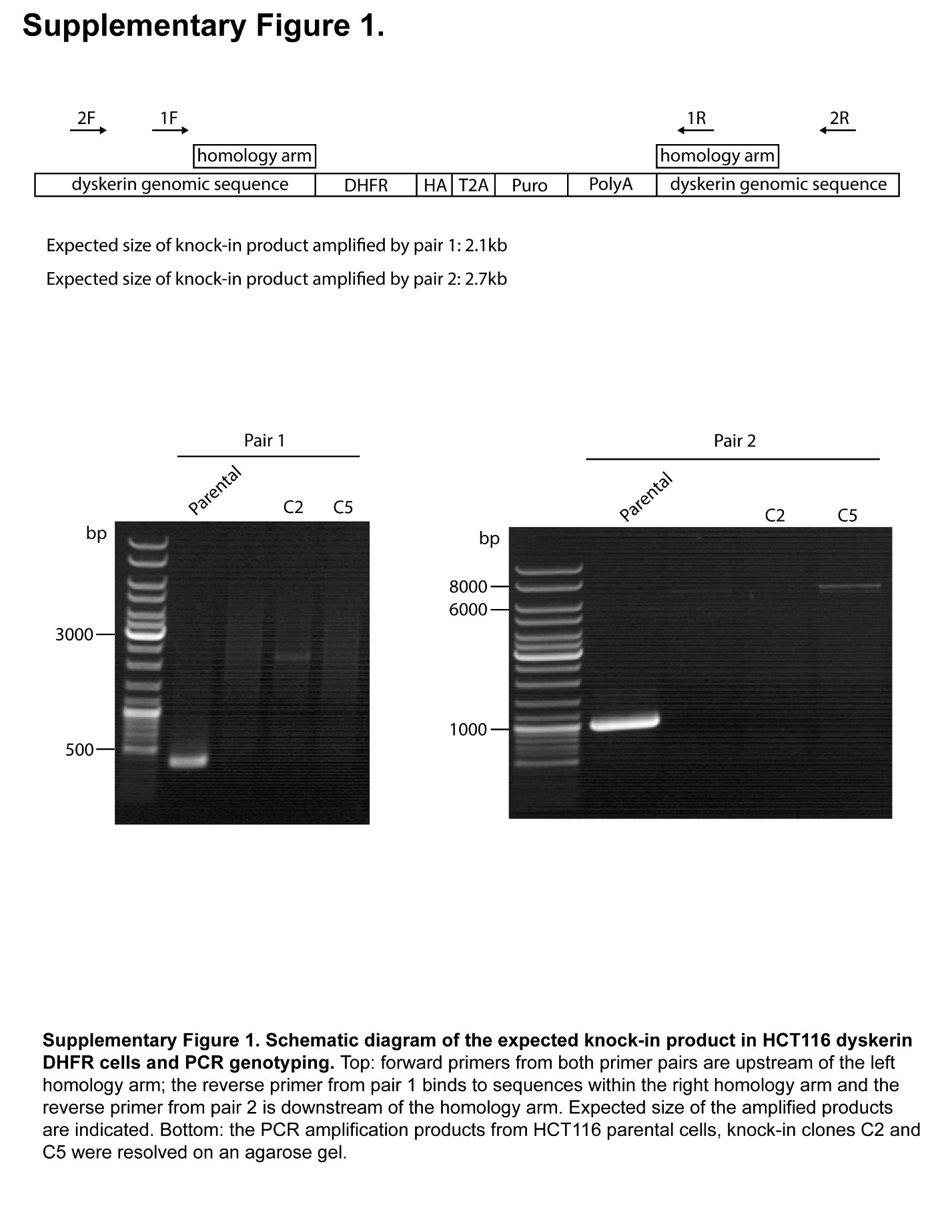

**Supplementary Figure 1. Schematic diagram of the expected knock-in product in HCT116 dyskerin DHFR cells and PCR genotyping.** Top: forward primers from both primer pairs are upstream of the left homology arm; the reverse primer from pair 1 binds to sequences within the right homology arm and the reverse primer from pair 2 is downstream of the homology arm. Expected size of the amplified products are indicated. Bottom: the PCR amplification products from HCT116 parental cells, knock-in clones C2 and C5 were resolved on an agarose gel.

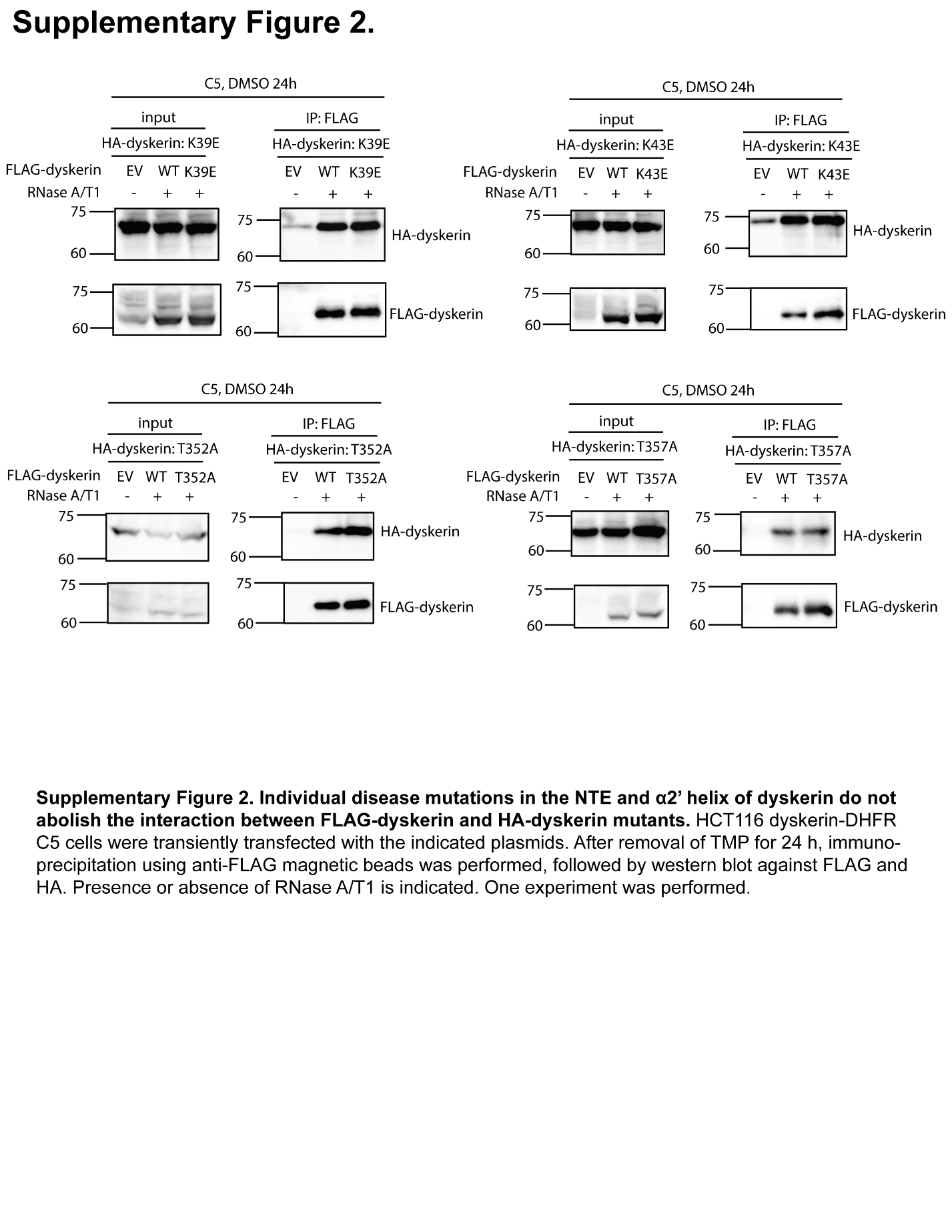

**Supplementary Figure 2. Individual disease mutations in the NTE and α2’ helix of dyskerin do not abolish the interaction between FLAG-dyskerin and HA-dyskerin mutants.** HCT116 dyskerin-DHFR C5 cells were transiently transfected with the indicated plasmids. After removal of TMP for 24 h, immunoprecipitation using anti-FLAG magnetic beads was performed, followed by western blot against FLAG and HA. Presence or absence of RNase A/T1 is indicated. One experiment was performed.

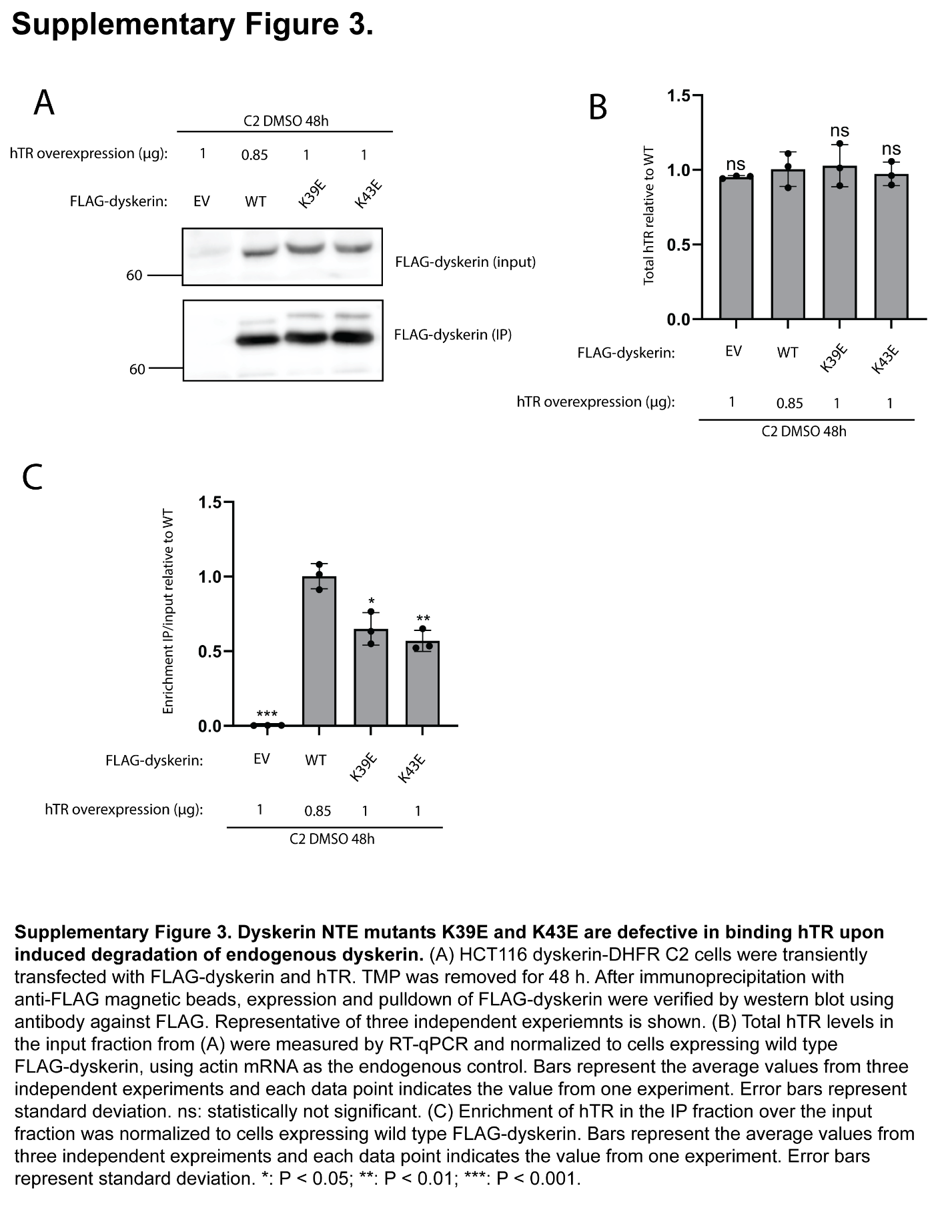

**Supplementary Figure 3. Dyskerin NTE mutants K39E and K43E are defective in binding hTR upon induced degradation of endogenous dyskerin.** (A) HCT116 dyskerin-DHFR C2 cells cultured in 10-cm plates were transiently transfected with FLAG-dyskerin and hTR. TMP was removed for 48 h. After immunoprecipitation with anti-FLAG magnetic beads, expression and pulldown of FLAG-dyskerin were verified by western blot using antibody against FLAG. Representative of three independent experiemnts is shown. (B) Total hTR levels in the input fraction from (A) were measured by RT-qPCR and normalized to cells expressing wild type FLAG-dyskerin, using actin mRNA as the endogenous control. Bars represent the average values from three independent experiments, and each data point indicates the value from one experiment. Error bars represent standard deviation. ns: statistically not significant. (C) Enrichment of hTR in the IP fraction over the input fraction was normalized to cells expressing wild type FLAG-dyskerin. Bars represent the average values from three independent experiments and each data point indicates the value from one experiment. Error bars represent standard deviation. *: P < 0.05; **: P < 0.01; ***: P < 0.001.

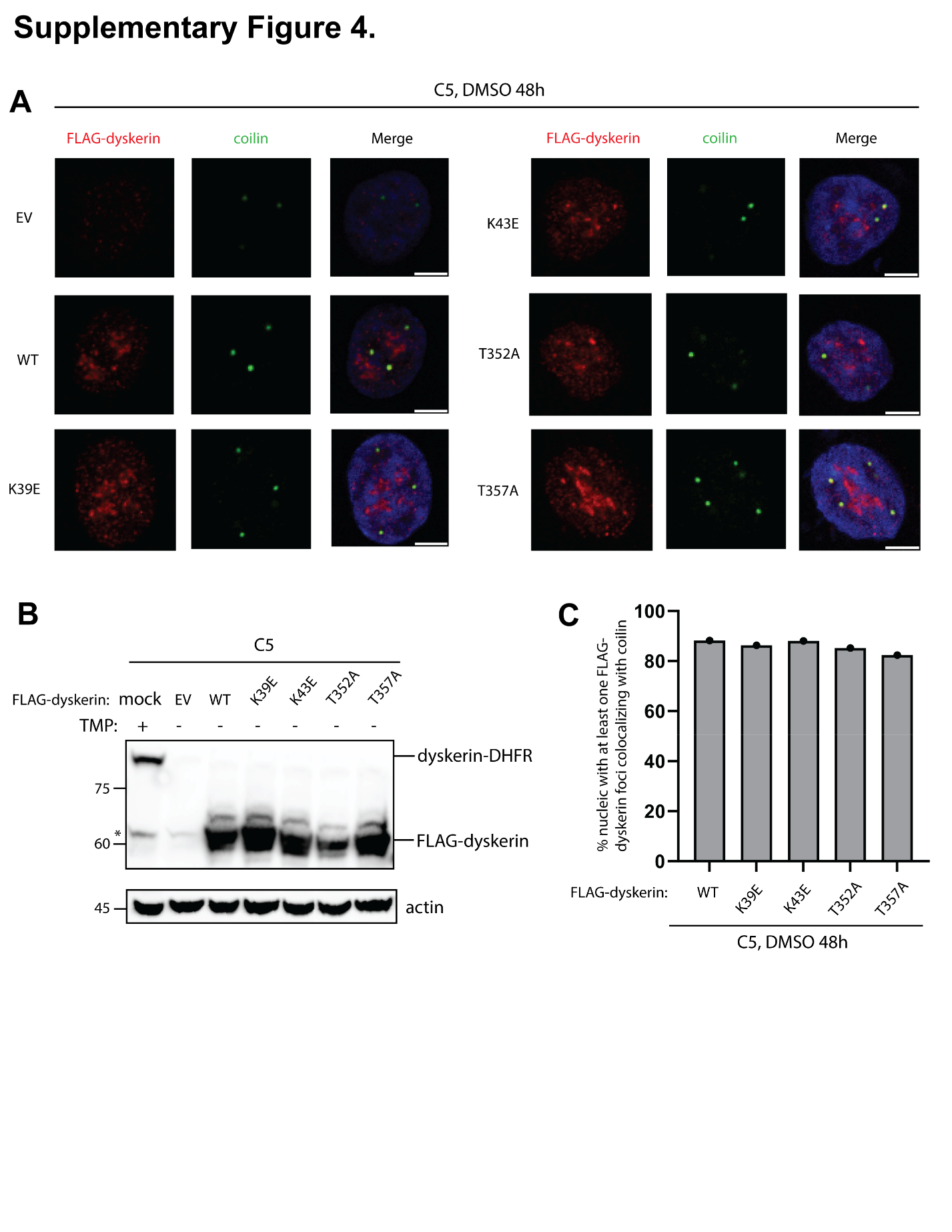

**Supplementary Figure 4. Disease mutations in the NTE and α2’ helix of dyskerin do not disrupt localization of dyskerin to CBs.** HCT116 dyskerin-DHFR C5 cells cultured in 6-cm plates were transiently transfected with 6 µg of EV/FLAG-dyskerin in TMP-free media for 6 h. Transfection mixture was then removed, and the cells were cultured in TMP-free media for another 42 h. (A) Dual indirect immunofluorescence was performed using primary antibodies against FLAG and the CB marker coilin. Nuclei were stained by DAPI. Scale bars indicate 5 µm. (B) Degradation of dyskerin-DHFR and expression of FLAG dyskerin was confirmed by western blot. Asterisk indicates non-specific band. (C) Quantification of the percentage of nuclei with at least one FLAG-dyskerin foci co-localizing with coilin. Bars represent the value from one experiment. At least 50 nuclei positive for both FLAG and coilin staining were examined in each sample.

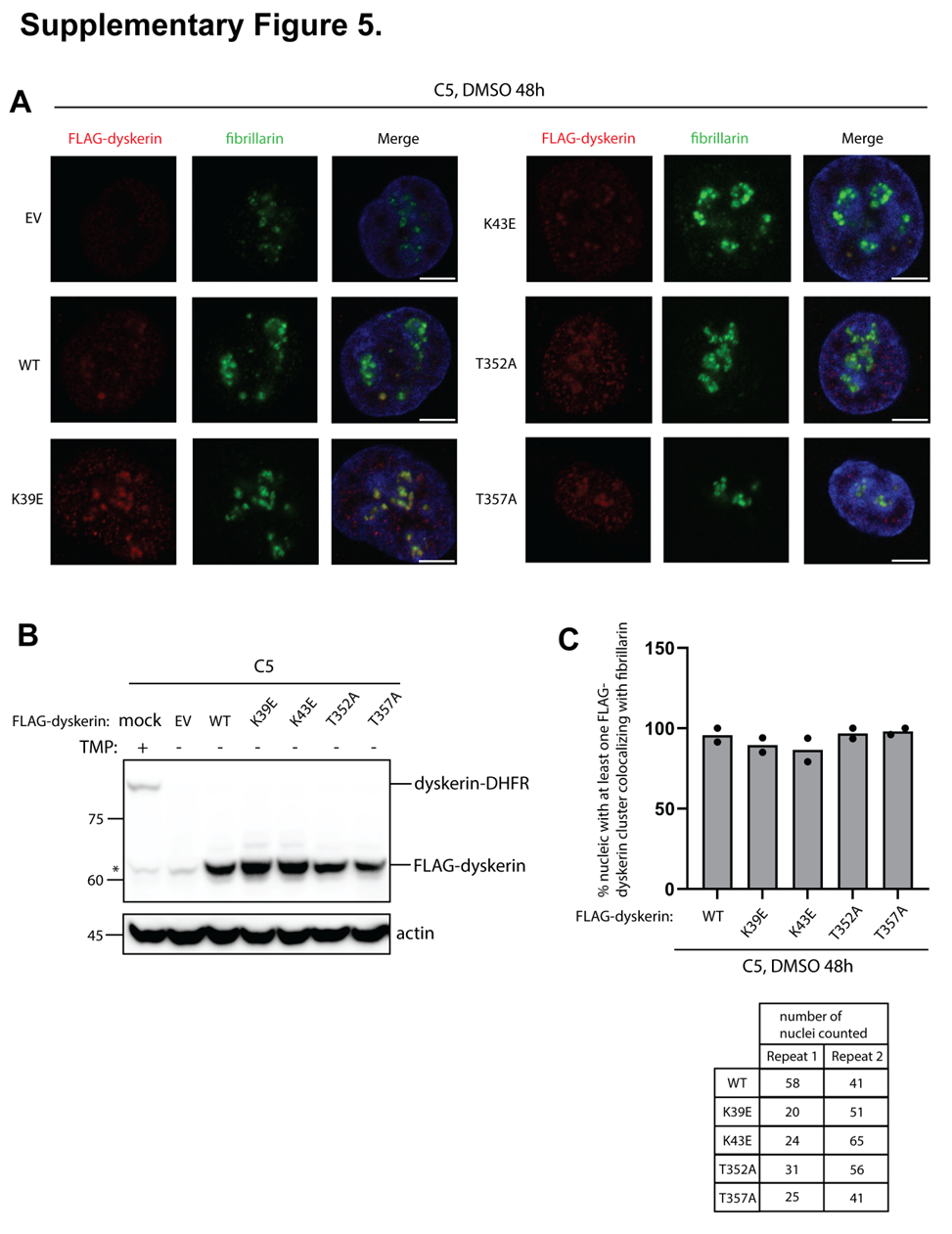

**Supplementary Figure 5. Disease mutations in the NTE and α2’ helix of dyskerin do not disrupt localization of dyskerin to the nucleolus.** HCT116 dyskerin-DHFR C5 cells cultured in 6-cm plates were transiently transfected with 6 µg of EV/FLAG-dyskerin in TMP-free media for 6 h. Transfection mixture was then removed, and the cells were cultured in TMP-free media for another 42 h. (A) Dual indirect immunofluorescence was performed using primary antibodies against FLAG and the nucleolar marker fibrillarin. Nuclei were stained by DAPI. Scale bars indicate 5 µm. (B) Degradation of dyskerin-DHFR and expression of FLAG dyskerin was confirmed by western blot. Asterisk indicates non-specific band. (C) Quantification of the percentage of nuclei with at least one FLAG-dyskerin clustering co-localizing with fibrillarin. Bars represent the average values from two independent experiments; each data point indicates the value from one experiment. The number of nuclei examined, positive for both FLAG and fibrillarin staining, were listed in the table.

**Supplementary Table 1. List of DNA oligos, siRNAs, and sgRNA**

| Name | Sequence | Usage |
| --- | --- | --- |
| hTR | F: 5’- CGGGTCTCTCGGGGGCGAGGGCGA-3’ | qPCR |
|  | R: 5’- GGTGCACGTCCCACAGCTCA-3’ |  |
| actin | F: 5’-TGAAGTGTGACGTGGACATC-3’ | qPCR |
|  | R: 5’-GGAGGAGCAATGATCTTGAT-3’ |  |
| TS | 5’-AATCCGTCGAGCAGAGTT-3’ | qTRAP |
| ACX | 5’-GCGCGGCTTACCCTTACCCTTACCCTAACC-3’ | qTRAP |
| 1F | 5’- CTCTACCACTTATACTTAGAGC-3’ | PCR genotyping |
| 1R | 5’- TCAATAAGGCTTTAGTTTCTCC-3’ | PCR genotyping |
| 2F | 5’-GAAAAGAAGAAGAGTAAGAAGG-3’ | PCR genotyping |
| 2R | 5’- CAGAGGGGATTTAAAAAATGC-3’ | PCR genotyping |
| DKC1 left homology arm | 5’- GGTATCTGTGAGCTTTCATTCTCTTTCTTTCTAGGACAGTGATACCACCAAGAAGAAGAAGAAGAAGAAGAAAGCAAAAGAGGTAGAATTGGTTTCTGAGGGAGGCGGTTACCCATAC-3’ (2 phosphorothioate bonds in the 5' end) | PCR amplify DHFR-HA-T2A-Puro-SV40 poly(A) |
| DKC1 right homology arm | 5’- TCTAGGACCTATGTTCCACTCTCTCAGCAACAAAAGGATCTATAACATGTTTTCTCAATAAGGCTTTAGTTTCTCCTCCAGCTTCAAGTGGCCTTCACTAGTCGACTGATCATAATCAGC-3’ (2 phosphorothioate bonds in the 5' end) | PCR amplify DHFR-HA-T2A-Puro-SV40 poly(A) |
